# The effect of ULV-based mosquito control on target and non-target organisms in Hungary: an experimental field study

**DOI:** 10.64898/2026.03.11.711007

**Authors:** László Zsolt Garamszegi, Gergely Nagy, Ágnes Klein, Tamara Szentiványi, Zsóka Vásárhelyi, Gábor Markó, Sándor Zsebők, Zoltán Soltész

## Abstract

Ultra-low volume (ULV) insecticide spraying with deltamethrin as the active ingredient is widely used in mosquito control programs, yet its effectiveness against target mosquitoes and its ecological side effects remain poorly quantified under field conditions in Central Europe. Here, we experimentally evaluated the short-term impact of ground ULV spraying on both mosquito populations and non-target flying insects in Hungary using a paired before–after-control–impact (BACI) design. Mosquitoes were sampled with BG Sentinel traps, while non-target insects were collected using malaise traps. ULV treatment resulted in a significant reduction in mosquito abundance at treated sites, with an average decline of approximately 45%. Native and invasive mosquito species, including *Aedes albopictus* and *Aedes koreicus*, showed similar proportional decreases. However, treatment effectiveness varied substantially among sites and was influenced by initial mosquito abundance and wind conditions. In parallel, malaise trap samples revealed a marked decline in non-target flying insects, with reductions exceeding 40% across multiple taxonomic groups, particularly among small- and medium-sized insects, and also when considering pollinator taxa together. Our results indicate that while ULV spraying can temporarily reduce mosquito abundance, it also imposes considerable short-term impacts on non-target insect communities, highlighting trade-offs between vector control and insect conservation within mosquito management programs.

## Introduction

Mosquitoes (Diptera: Culicidae) are globally recognized as major vectors of pathogens affecting both humans and animals, transmitting pathogens that pose significant public health burdens (Becker et al. 2020). Their control has therefore become an essential component of disease prevention and nuisance reduction programs worldwide. The current paradigm for sustainable mosquito suppression is Integrated Vector Management (IVM), which promotes the use of multiple, complementary approaches tailored to local ecological and epidemiological conditions (van den Berg & Takken 2009; WHO 2012). Core elements of IVM include environmental management and habitat modification, biological control, targeted use of larvicides, and, where appropriate, adulticidal interventions, combined with community engagement and the consideration of insecticide resistance (Martinou et al. 2020).

Despite the recognized importance of integrated approaches, in many countries mosquito control efforts remain heavily reliant on space spraying of insecticides (typically pyrethroid-based formulations like deltamethrin) against adult mosquitoes, most often through ultra-low volume (ULV) applications from ground vehicles (Bonds 2012). These treatments are widely perceived as an efficient and rapid method to reduce mosquito abundance and to provide visible relief from biting populations (Baldacchino et al. 2015). Nevertheless, it may affect different mosquito species differently, depending on their daytime activity patterns, their degree of susceptibility, and the application method (Boubidi et al. 2016, Walker et al. 2025). This variability in response to the treatment is particularly concerning in the context of ongoing shifts in mosquito species composition worldwide.

The threat mosquitoes pose started to increase further in recent decades, as the global distribution of invasive mosquito species has expanded rapidly, facilitated by international trade, human mobility, and changing climatic conditions (Garcia-Rejon et al. 2021; Kraemer et al. 2015; Medlock et al. 2015; Murray et al. 2013). Species such as *Aedes albopictus* (the Asian tiger mosquito) and *Aedes aegypti* (the Egyptian mosquito) have colonized large areas outside their native ranges, establishing stable populations in many temperate regions. These invasive taxa are of particular concern because they are competent vectors of a range of arboviruses, including dengue, chikungunya, zika, and West Nile virus, and are therefore increasingly regarded as priority targets for mosquito control programs (Leta et al. 2018).

These invasive mosquitoes present unique challenges for conventional control approaches (Baldacchino et al. 2015; Corcos et al. 2020; Farajollahi et al. 2012; Fonseca et al. 2013; Stiles et al. 2025), as many widely implemented control measures are considered suboptimal for invasive *Aedes* mosquitoes (Gubler 1998; Reiter & Gubler 1997). One reason for this might be that the daytime activity of these species can be hypothesized to reduce the effectiveness of ULV applications, which are typically conducted during evening or night-time hours. Moreover, their preference for resting in sheltered urban microhabitats – such as areas around buildings, fences, and dense vegetation – can also limit their exposure to airborne insecticide droplets.

Three invasive mosquito species are currently present in Hungary (Garamszegi et al. 2025; Sáringer-Kenyeres et al. 2019). The establishment and spread of *Ae. albopictus*, *Ae. japonicus*, and *Ae. koreicus* raise increasing public health risks as their presence may facilitate the introduction or local transmission of various arboviruses, while the tiger mosquito contributes to considerable nuisance levels during the summer months – in addition to native *Culex*, *Aedes* and *Culiseta* mosquitoes. Mosquito control in Hungary is organized through a centrally coordinated national program, which prioritizes large-scale, rapid reduction of adult mosquito populations (Csiba et al. 2025; Sáringer-Kenyeres 2020). This program relies overwhelmingly on the use of ULV insecticide applications carried out from ground vehicles, which replaced aerial spraying methods that dominated in the past. Although biological control methods, like the one based on the use of *Bacillus thuringiensis israelensis* gains interest in the society, this approach is still heavily underrepresented in practice. Although the ULV spraying of insecticides is widely perceived as effective in reducing immediate nuisance, its efficiency against both native and invasive mosquito species remains questionable, while the dominance of this approach raises concerns about its sustainability and broader ecological consequences (Meftaul et al. 2020; Schleier & Peterson 2013). The ecological consequences of the applied practice have rarely been evaluated under field conditions in Hungary, or were not done along scientific standards (Pénzes 1995; Sáringer 1988; Török et al. 2020). This lack of empirical data is especially problematic given the behavioural and ecological characteristics of invasive *Aedes* species, which make them less likely to be affected by conventional spraying operations. Consequently, it is unclear to what extent the current strategy achieves its stated objectives of reducing the abundance of both native and invasive mosquito populations.

To quantify the efficiency of the local ULV spraying method on killing the target mosquitoes and also to characterise its ecological side effects in Hungary, we designed an experimental sampling regime (before–after-control–impact, BACI) around the particular events of the mosquito control. We selected both treatment and control sites based on their involvement in the centrally organized mosquito control program, and sampled the flying insect fauna both before and after the application of insecticides by using different traps that preferentially capture different groups of insects. We predicted that if the ULV spraying was effective against the mosquitoes (target group), then the number of mosquitoes would have decreased in the treatment group after the dispersion of chemicals, but not in the control sites.

Furthermore, along the hypothesis that this method is less effective against invasive than native mosquitoes, we expected to observe effects with different magnitudes when the data are split between native and invasive mosquito species. Finally, if the insecticide treatment has an unwanted ecological impact on non-target organisms, we predicted to find fewer flying insects in general in the traps after the treatment.

## Materials and Methods

### Study sites

Three companies in Hungary were contacted to obtain publicly available information about their organised mosquito control program in 2024 and 2025 (i.e., planned dates and prospective routes of the spraying vehicle). In 2024, our research team performed a study focusing on the mosquito control program in one residential area of the capital (Budapest, 10^th^ district, 4 treatment sites, Figures 1A and 1B). We selected three localities in the agglomeration area (Dunakeszi, Telki, Veresegyház, 4 control sites, Figures 1A), where there was no ULV treatment performed at the same time, and we could logistically arrange simultaneous trapping of mosquitoes and other insects (see further details below).

**Figure 1.**
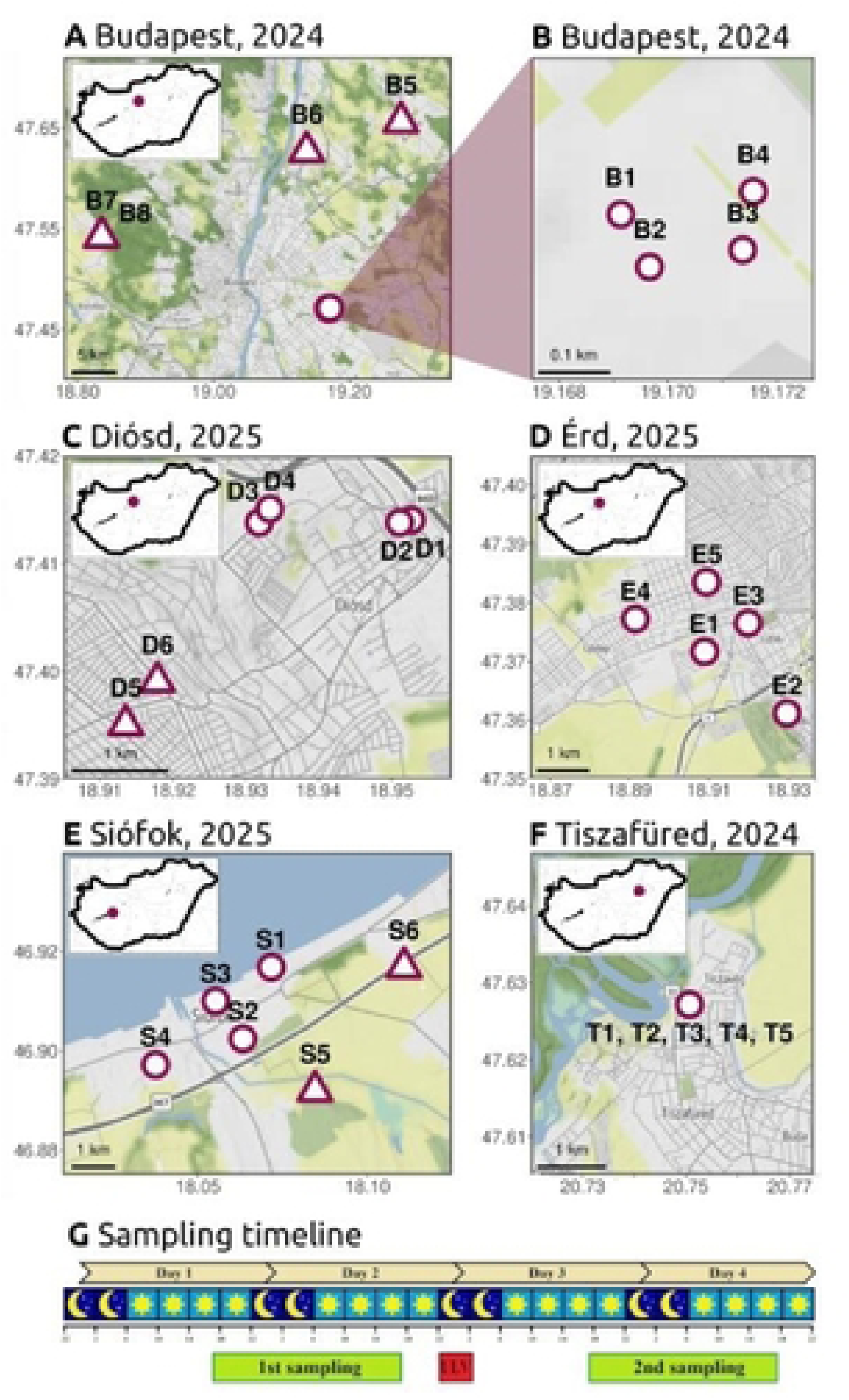
Sampling sites and sampling regime used to investigate the effect of ULV ground treatment of insecticides on the abundance of target and non-target organisms in Hungary. Maps (A-F) show the locations of each sampling sites in different municipalities and years. Circles are for treatment sites, triangles are for control sites. The timeline (G) gives the temporal organisation of sampling sessions before and after the ULV treatment, which was applied in parallel in the treatment and control sites.

In 2025, to improve sampling efficiency and to extend the study to other distant localities, we organized sample collection with the involvement of the public. Based on the known dates and routes of the control program, at three localities (Érd, Diósd, Siófok, Figures 1C - 1E), we recruited volunteers. A targeted social media campaign was performed, in which the aims of the study and the expected contributions were appropriately explained. This was necessary, because we particularly sought sampling sites at the close proximity of the route of the ULV ground vehicle, where we could safely install different traps in private properties, and we could also rely on the help of volunteers to collect the trap materials in concert by using the same standards that we applied in 2024. Interested citizens could sign up for hosting the traps through an online application form, where they also gave informed consent to be part of the study and agreed to the handling of their personal data. Based on the level of engagement, we could select multiple sites in each location. The only criteria for involvement were that the volunteers’ property was close enough (< 100 m) to the prospected route of the ULV vehicle, and the owners were available for emptying the traps around the suspected date of treatment. In the same way, we also selected volunteers for the control sites, ensuring that the control points were located within a 3 km radius of the treated area and at least 1 km from the ULV vehicle route. In Érd, we initially defined control sites, but later these points were considered as treated sites due to a change in the realized ULV vehicle route with regard to the planned route. Altogether, we operated traps at 13 treatment and 4 control sites by the help of volunteers in 2025 in the three localities (Figures 1C - 1E). Each volunteer was given an appropriate training for the operation of traps and also about the required preparation steps of the insects collected (personal demonstration, printed guide). The traps were installed and set in operation in the sites by professionals, volunteers were only asked to collect the caught insects at given dates and times and store them. Volunteers were compensated for their efforts with a gift package, and later received information about the species observed in their gardens. Given that for the successful recruitment it was necessary to inform the volunteers about the purpose of the study, and also that the mosquito control program is advertised in each municipality, it was not possible to assign volunteers blindly with respect to the treatments. Therefore, we can only assume that having information on the mosquito control program did not affect the volunteers’ behaviour when emptying the traps, and this did not raise bias on our data. The companies that performed the insecticide treatment were not informed about the exact location of the sampling sites. Given that the sampling regime were different in 2024 and 2025 with regard to distance between the treatment and control sites, we considered this potentially confounding effect in the level of statistical analysis (see the mixed modelling approach below and Tables S1-S3).

In 2025, in one locality (Tiszafüred, 1 site sampled at 5 different occasions around the dates of the realized insecticide treatment, Figure 1F), the traps were installed and operated along our established standards by a company who performed the mosquito control program. The collected samples were then transferred to our laboratory for further processing for the purpose of the study. However, the use of the data for this locality could raise issues about non-independence, thus we repeated our statistical analyses with and without catch results from Tiszafüred. Given that these parallel tests yielded remarkably similar conclusions, in the main text we present the results that correspond to the larger sample size, but in Table S4 we also show the main outcomes without considering the multiple samples from this locality.

### ULV application details

Ground ULV interventions occurred between 10:00 pm and 02:00 am, and were performed by professional companies with the appropriate licences and in compliance with the applicable legislation and technological regulations. The spraying of Deltasect Plus 20 ULV (2.22% deltamethrin + 0.22% *Chrysanthemum cinerariaefolium,* distributor: Sharda Hungary Ltd., permission nr: 30430/2022/KBKHF) with 1:15 dilution or Deltasect Plus 1.2 ULV (0.135% deltamethrin + 0.013% *Chrysanthemum cinerariaefolium*, distributor: Sharda Hungary Ltd., permission nr: 47799-5/2020/JIF) without dilution was performed using ground-based truck-mounted mist sprayers. The dose for the emitted products was calibrated to reach 0.6-0.8 liter/ha leading to deltamethrin concentration of c.a. 1 gram/ha. The vehicle aimed to follow the pre-defined routes with a maximum speed of 20 km/h.

### Sample collection and taxonomic identification

In each site, we aimed to install two types of traps (one of each), in close proximity to each other in the same garden. BG Sentinel traps were used to sample the mosquito community (Medlock et al. 2018), and these were supplied with CO_2_ and lure as attractants. The BG Sentinel traps were c.a. 25 m away from the route of the spraying vehicle (mean: 25.23 m, range: 3.02-75.4 m). Malaise traps were operated to catch a broad spectrum of species of flying insects with minimal bias (Uhler et al. 2022; see Báldi et al. 2022 for pictures on malaise traps used in this study), and these were installed with a c.a. 22 m distance from the route of the vehicle (mean: 21.78 m, range: 3.62-75.78 m).

At each site, we aimed to apply the following sampling regime in both the control and treatment sites in parallel (Figure 1G). The first sampling started a day before the planned ULV treatment, when the traps were operated for c.a. 24 hours and captured insect communities still unaffected by the application of insecticides. The same sampling window was used from the next day, after the event of ULV spraying in the given municipality for the treatment site. In few instances, the mosquito control program was delayed due to bad weather conditions. In such cases the second sampling window was also shifted in order to sample a 24-hour period starting from the afternoon right after the night of the treatment.

The content of the BG Sentinel traps was processed by a taxonomist expert (Z.S.) under a microscope. Specimens for all mosquitoes were identified at the species level on morphological characteristics (Briët et al. 2021; Kenyeres & Tóth. 2008). Individuals could have been sexed based on their morphology, but for the current study, counts for females and males were combined (91.1% of the trapped individuals were females). During the identification procedure, the list of species and their respective abundances (i.e., the number of individuals) were determined for each BG Sentinel trap (see the list of species in Table 1). However, for the purpose of the study, we have not analysed the effect of insecticide treatment for each mosquito species separately, but followed the change in abundance for all mosquitoes combined. In addition, we also separated native and invasive species and tested for treatment effects on abundance separately in these two groups.

**Table 1.** The number of individuals of mosquito species trapped in 2024 and 2025 to test the effect of ULV treatment on the abundance of mosquitoes. The scientific names of mosquitoes are written based on Snow and Ramsdale (2003). Data are given separately for treatment and control sites and for before and after treatment statuses.

|  | TREATMENT SITES |  | CONTROL SITES |  |
| --- | --- | --- | --- | --- |
|  | before | after | before | after |
| *Aedes albopictus* | 369 | 153 | 113 | 76 |
| *Aedes koreicus* | 3 | 1 | 1 | 1 |
| *Aedes sticticus* | 80 | 34 | 0 | 0 |
| *Aedes vexans* | 179 | 37 | 1 | 0 |
| *Anopheles maculipennis* | 3 | 3 | 0 | 2 |
| *Anopheles hyrcanus* | 11 | 0 | 0 | 0 |
| *Coquillettidia richiardii* | 72 | 23 | 6 | 1 |
| *Culex hortensis* | 1 | 1 | 1 | 0 |
| *Culex pipiens* | 485 | 395 | 259 | 165 |
| *Culiseta annulata* | 0 | 0 | 4 | 2 |
| *Ochlerotatus annulipes* | 17 | 7 | 0 | 0 |
| *Ochlerotatus caspius* | 0 | 0 | 1 | 1 |
| **TOTAL** | **1220** | **654** | **386** | **248** |

Given the large number of individuals and the diversity of insect groups caught, the content of the malaise traps was processed at the order level, i.e., the number of individuals falling into the major taxonomic groups were estimated. In particular, the order of Diptera (without mosquitoes), Hymenoptera, Hemiptera, Lepidoptera, and Coleoptera were represented by a considerable number of individuals, while the remaining insects were pooled into the “other insects” category. Beyond these taxonomic groupings, we also defined functional categories. One of these relied on the size of the individuals given that the applied methodology for mosquito control supposedly affects flying insects with a size similar or smaller than mosquitoes (Boyce et al. 2007; Kwan et al. 2009). Accordingly, we also pooled individuals based on their body size and defined three categories: i) insects smaller than mosquitoes (< 3 mm), ii) insects more or less with the size of mosquitoes (3-10 mm), and iii) insects larger than mosquitoes (> 10 mm) (Mihályi & Sztankayné Gulyás 1963). The other functional category was created for pollinators that included all individuals from the following taxonomic groups: Syrphidae, Apidae and Lepidoptera.

### Environmental variables

We also investigated whether certain variables reflecting the environmental conditions of the insecticide treatment might affect its efficiency in terms of the degree by which it decreases the abundance of mosquitoes (or our efficiency to catch mosquitoes with the BG Sentinel traps). For this, we considered the following variables with potential effect: i) the shortest distance between the geographic position of the BG Sentinel trap and the path of the ULV vehicle; ii) the time, relative to the sunset, of the vehicle passing by the property where the traps were installed; and iii) climatic conditions defined by temperature, precipitation and wind speed. Information on the realized route and timing of the ULV vehicle for calculating i) and ii) was available in the log file of GPS logger with which the ULV vehicle should be equipped by law (this information was extracted after the event of the insecticide treatment). Georeferenced weather data for iii) were gathered for each sampling location using the Hungarian Meteorological Service (HungaroMet) open Meteorological Database (odp.met.hu). We obtained location-specific temperature (daily average, in °C with 0.01 accuracy) and precipitation (daily sum, in mm with 0.01 accuracy) and maximum wind speed (km/hour) for each 24-hour trapping session. There is a strong correlation between mean and maximum wind speed using a reference data (Pearson’s *r* = 0.603, N = 156, P < 0.001).

### Data analyses

The distribution of variables was checked before analyses and the appropriate transformation was applied if it was necessary to reach assumptions about normal distribution. Importantly, for the analyses focusing on the abundance of particular taxonomic groups, we systematically applied log_10_-transformation on the number of individuals (for illustrative purposes we show the original data on the graphs).

The focal analysis relied on a paired design, in which the abundance of particular insect taxa was compared at each site before and after the application of insecticide treatment (i.e., each site is its own control). Accordingly, we performed paired t-tests separately for the treatment and control sites, and investigated if the number of individuals at the post-treatment conditions were different than at the pre-treatment conditions. These paired t-tests (both for the treatment and control sites) were performed for each target and non-target group considered, and the corresponding *P* values are shown on the figures.

Given that the data were hierarchically structured (more than one site from the same location) and that were not randomly varied with respect to date and year (note the difference in the sampling design between 2024 and 2025), we also tested our predictions by using mixed models that allow accounting for several potential confounding factors. In these models, the response variable was the abundance of the focal taxa, the fixed predictors were status (before or after treatment), year (as a categorical variable), and date (ordinal date), while the random part included effects for locality and sites nested within locality. The significance of the fixed terms was determined by likelihood ratio tests, in which we compared the deviance of the full model with that of a reduced model that lacked the term. These models were fit separately for the treatment and control sites, and shown in the Supplementary Material (Tables S1-S3).

The efficiency of the insecticide treatment (or our capturing sessions) was assessed by focusing on the difference between the number of mosquito individuals after and before the treatment at sites where ULV spraying was performed. When the abundance of mosquitoes declined considerably (i.e., large difference between the pre- and post-treatment conditions), the applied mosquito control method could be considered very successful. Hence, a large negative difference indicated high efficiency. There were two sites, where the number of mosquitoes was actually higher after than before the treatment. We did not calculate differences with a positive sign at these sites (because it is unlikely that the abundance of mosquitoes increased due to the treatment), but we assigned 0 values for them to reflect minimum efficiency. We investigated the relationship between the efficiency of the treatment (difference in the number of mosquitoes) and the environmental variables one by one based on a correlation approach. Given that we performed multiple tests for the same hypothesis, we adjusted the *P* values by using false discovery rates (Benjamini & Hochberg 1995).

Data transformation, statistical analyses and the figures were done in the R statistical environment (R Core Team 2023). The mixed models were performed by using the package lme4 (Bates et al. 2015).

## Results

### Target organisms: mosquitoes (BG Sentinel traps)

Within the treatment group, there was a significant negative effect of insecticide application on the abundance of mosquitoes (Figure 2A). Before the treatment, the BG Sentinel traps captured 65.23 (SE = 10.84) individuals on average. The mean number changed to 34.73 (SE = 9.09) after the ground treatment, which corresponds to a 46.76% decline on average in the number of mosquito individuals trapped. In contrast, the same tendency was not observed in the control areas, where no mosquito control was performed (group means ± SE: before treatment, 21.38 ± 5.82; after treatment, 17.25 ± 4.25; Figure 2A). The mean number of mosquitoes was significantly higher in the treatment than in the control areas before the day of ground spraying at the treatment sites (*t* = 3.181, df = 11.236, *P* = 0.008). However, such a significant difference was not observable after the day of treatment (*t* = 1.295, df = 10.21, *P* = 0.224). These results were very similar when we performed mixed models that accounted for the non-independence of data, for the difference in sampling design between years, and also for the potentially confounding effect of sampling date and spatial aggregation (Supplementary Material Table S1).

**Figure 2.**
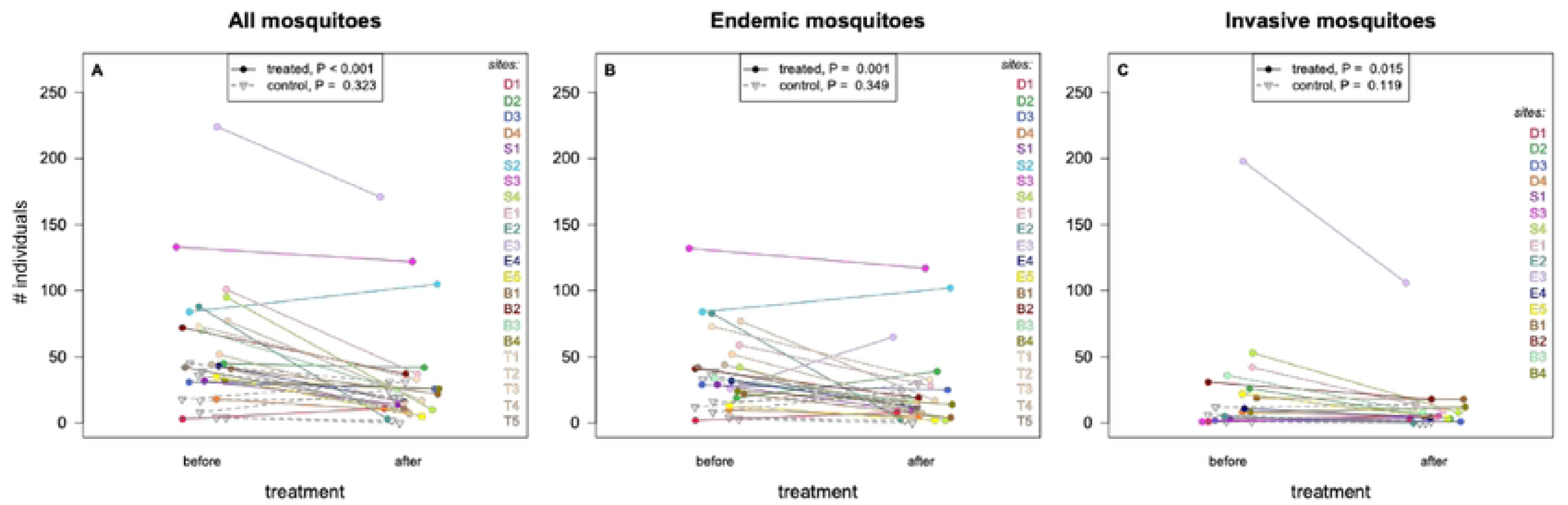
The effect of insecticide treatment on the abundance of mosquitoes (A: all species considered together, B and C: endemic and invasive mosquitoes treated separately). Coloured dots are for sites where mosquito control was applied, grey triangles are for sites where such intervention was not carried out at the same date. Different colours represent different sites (with IDs shown on the right corresponding to locations shown in Figure 1), and the lines connect the detected mosquito abundances of the same site before and after the night of the application of insecticides. *P* values are from the corresponding paired-t test performed on the log_10_-transformed data. Points along the x-axes are jittered for better visualisation.

When we separately analysed the native and invasive species, we found parallel results. The decline in mosquito abundance in the treatment group was also significant when individuals belonging to native species were considered only (group means ± SE: treatment group, before insecticide application, 44.05 ± 6.50; treatment group, after insecticide application, 25.18 ± 6.62; control group, before insecticide application, 18.88 ± 5.25; control group, after insecticide application, 14.62 ± 3.62; Figure 2B). The number of individuals of invasive species was also significantly lower in the after-treatment sample (group means ± SE: treatment group, before insecticide application, 29.12 ± 11.95; treatment group, after insecticide application, 12.94 ± 6.35; control group, before insecticide application, 5 ± 2.61; control group, after insecticide application, 3.75 ± 3.42; Figure 2C). The proportional change in the abundance was considerable for both native and invasive mosquitoes (42.83% and 55.58% decline on average, respectively).

We detected remarkable differences in how the ULV spraying left signatures on the number of captured mosquito individuals (Figure 2). We investigated if this variance could have been caused by the effect of some specific circumstantial variables. We found that the decline in abundance of all mosquitoes at particular treatment sites was significantly associated with the initial mosquito abundance (*r* = −0.582, N = 22, *P* = 0.005, Figure 3A) and maximum wind power detected during the sampling window (*r* = 0.590, N = 22, *P* = 0.004, Figure 3A), but not with the distance from the path of the ULV spraying vehicle (*r* = −0.310, N = 22, *P* = 0.160), the timing of the spraying (relative to the time of sunset, *r* = −0.029, N = 22, *P* = 0.896), precipitation (*r* = −0.096, N = 22, *P* = 0.672), and temperature (*r* = 0.412, N = 22, *P* = 0.057). The relationships for initial mosquito abundance and wind power were also significant when we controlled for the number of tests performed (both *P* = 0.014). Furthermore, when we held constant the effect of initial mosquito abundance in a multiple regression approach, the slope for wind speed remained significant (*P* = 0.027).

**Figure 3.**
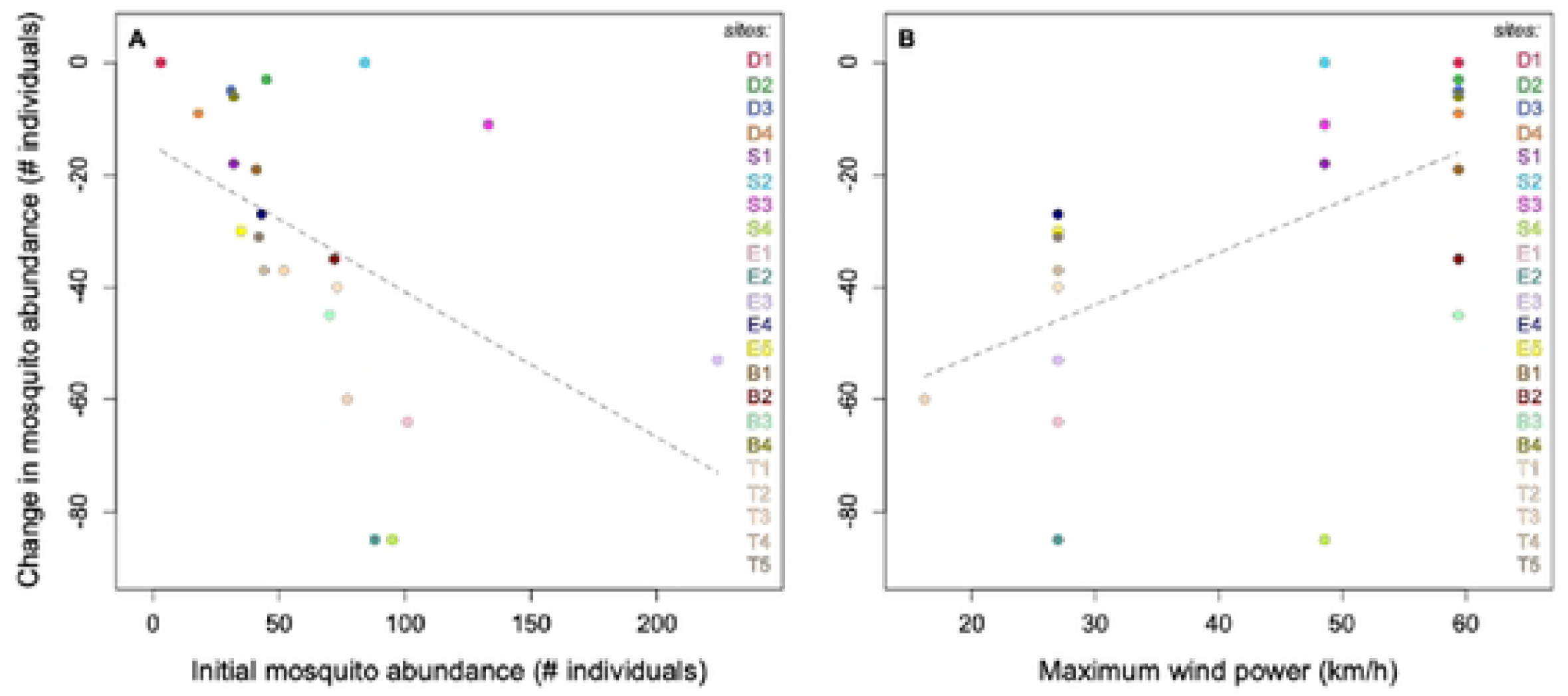
The relationship between initial mosquito abundance, maximum wind power and the effect of ULV treatment in terms of the decline in the number of mosquito individuals captured in the BG Sentinel traps. Different colours represent different sites (with IDs shown on the right corresponding to locations shown in Figure 1), grey lines are regression lines based on the raw data. Sites, where the number of mosquitoes increased after the ULV treatment (D1, S2) were considered with 0 values for the y-axis variable reflecting the minimum of the effectiveness of the treatment (see text for more details).

### Non-target organisms: other insects (Malaise traps)

In the malaise traps, we detected statistical evidence for treatment effect at the sites where ULV spraying was applied (Figure 4). The application of insecticides significantly reduced the abundance of non-target organisms at all but one treatment site (while such an effect was not detected for the control group). We further investigated whether the effect of ULV spraying detected in the malaise traps affected different insect groups dissimilarly or not.

**Figure 4.**
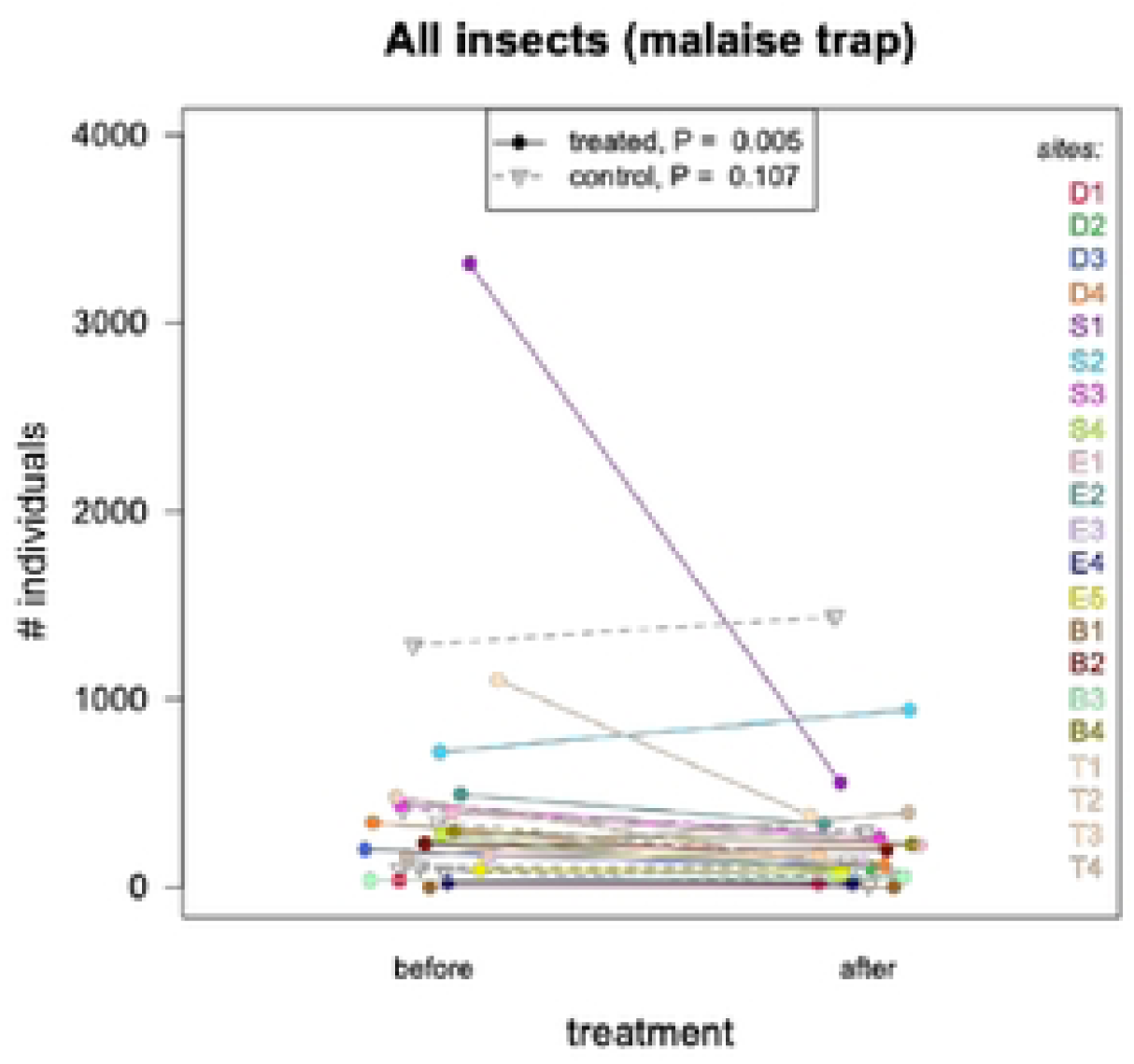
The effect of insecticide treatment on the abundance of non-target organisms as detected in the malaise traps. Coloured dots are for sites where mosquito control was applied, grey triangles are for sites where such intervention was not carried out on the same date. Different colours represent different sites (with IDs shown on the right corresponding to locations shown in Figure 1), and the lines connect the detected insect abundances of the same site before and after the night of the application of insecticides. *P* values are from the corresponding paired-t test performed on the log_10_-transformed data. Points along the x-axes are jittered for better visualisation.

First, we separated the captured individuals based on their size (i.e., small < smaller than mosquito; medium size ~size of a mosquito; large > larger than mosquito). We found that the significant decrease in the abundance of non-target organisms due to insecticide treatment was prevalent in the small and medium size insects but not in large insects (Figure 5).

**Figure 5.**
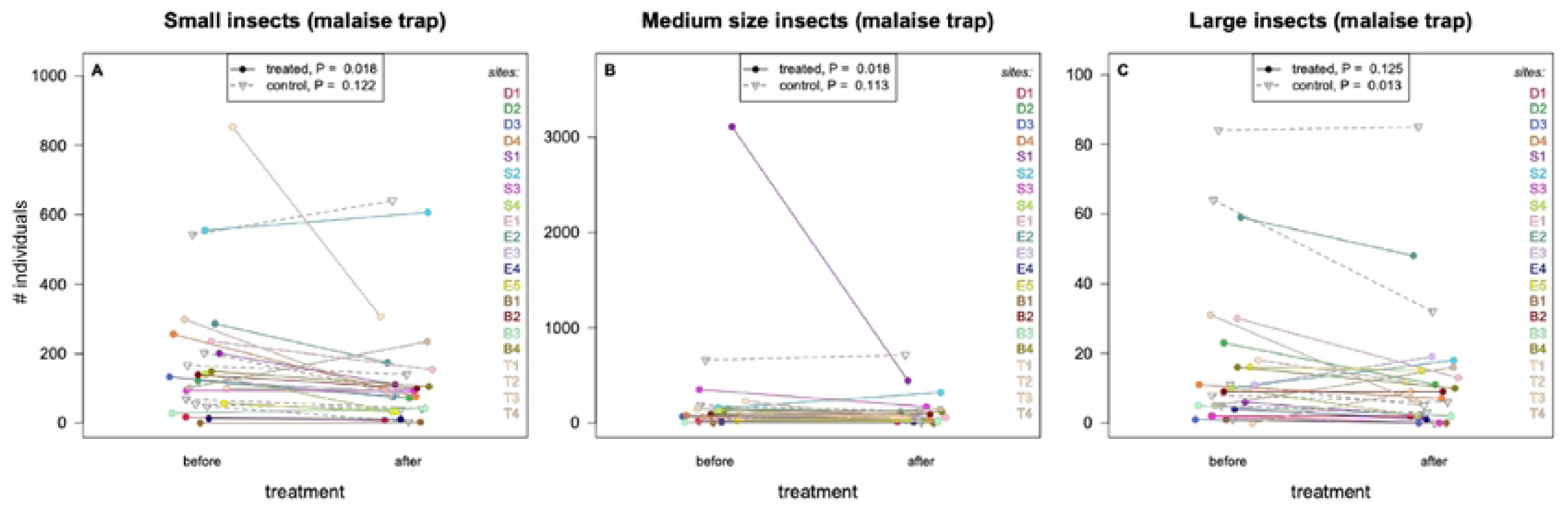
The effect of insecticide treatment on the abundance of non-target organisms in the malaise traps as detected in different size groups (A: insects smaller than mosquitoes, B: insects more or less with the size of mosquitoes, C: insects larger than mosquitoes). Coloured dots are for sites where mosquito control was applied, grey triangles are for sites where such intervention was not carried out on the same date. Different colours represent different sites (with IDs shown on the right corresponding to locations shown in Figure 1), and the lines connect the detected insect abundances of the same site before and after the night of the application of insecticides. *P* values are from the corresponding paired-t test performed on the log_10_-transformed data. Points along the x-axes are jittered for better visualisation.

Second, using the paired experimental design that we adopted above, we analysed the effect of mosquito control on the abundance of insects by tabulating data into taxonomic order (Figure 6). The negative trend was apparent in all of the major orders investigated, all of which was significant except the case for Hymenoptera. Focusing on the total number of individuals that were captured altogether at the treatment sites, we found a considerable reduction in the abundance of insects in each major insect order when compared with the control sites (Table 2). On average, at the treatment site the reduction of abundance was 41.33% while in the non-treated sites it was 12.32%.

**Figure 6.**
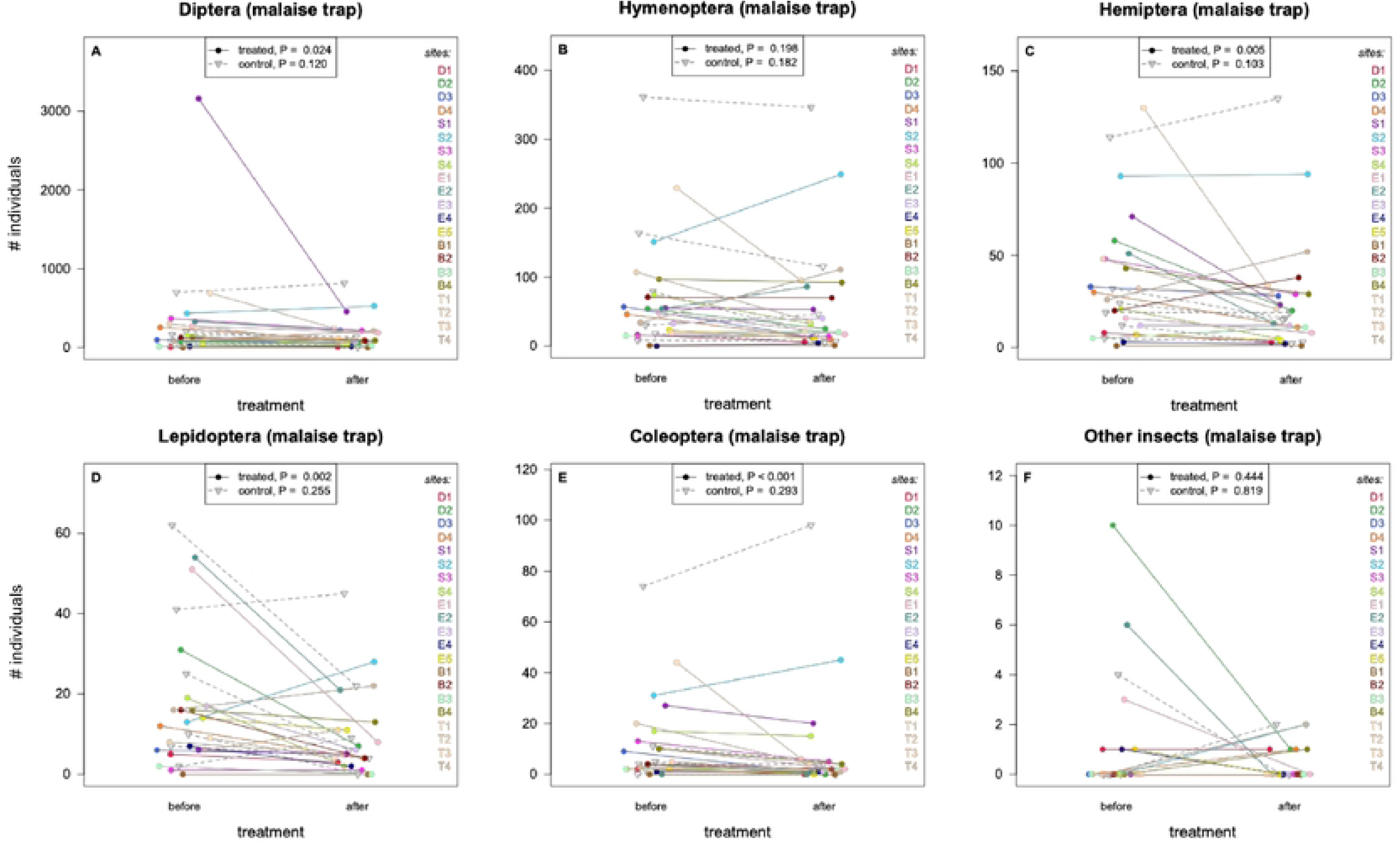
The effect of insecticide treatment on the abundance of non-target organisms in the malaise traps as detected in different taxonomic groups of insects (A: Diptera without mosquitoes, B: Hymenoptera, C: Hemiptera, D: Lepidoptera, E: Coleoptera, F: other insects). Coloured dots are for sites where mosquito control was applied, grey triangles are for sites where such intervention was not carried out on the same date. Different colours represent different sites (with IDs shown on the right corresponding to locations shown in Figure 1), and the lines connect the detected insect abundances of the same site before and after the night of the application of insecticides. *P* values are from the corresponding paired-t test performed on the log_10_-transformed data. Points along the x-axes are jittered for better visualisation.

**Table 2.**
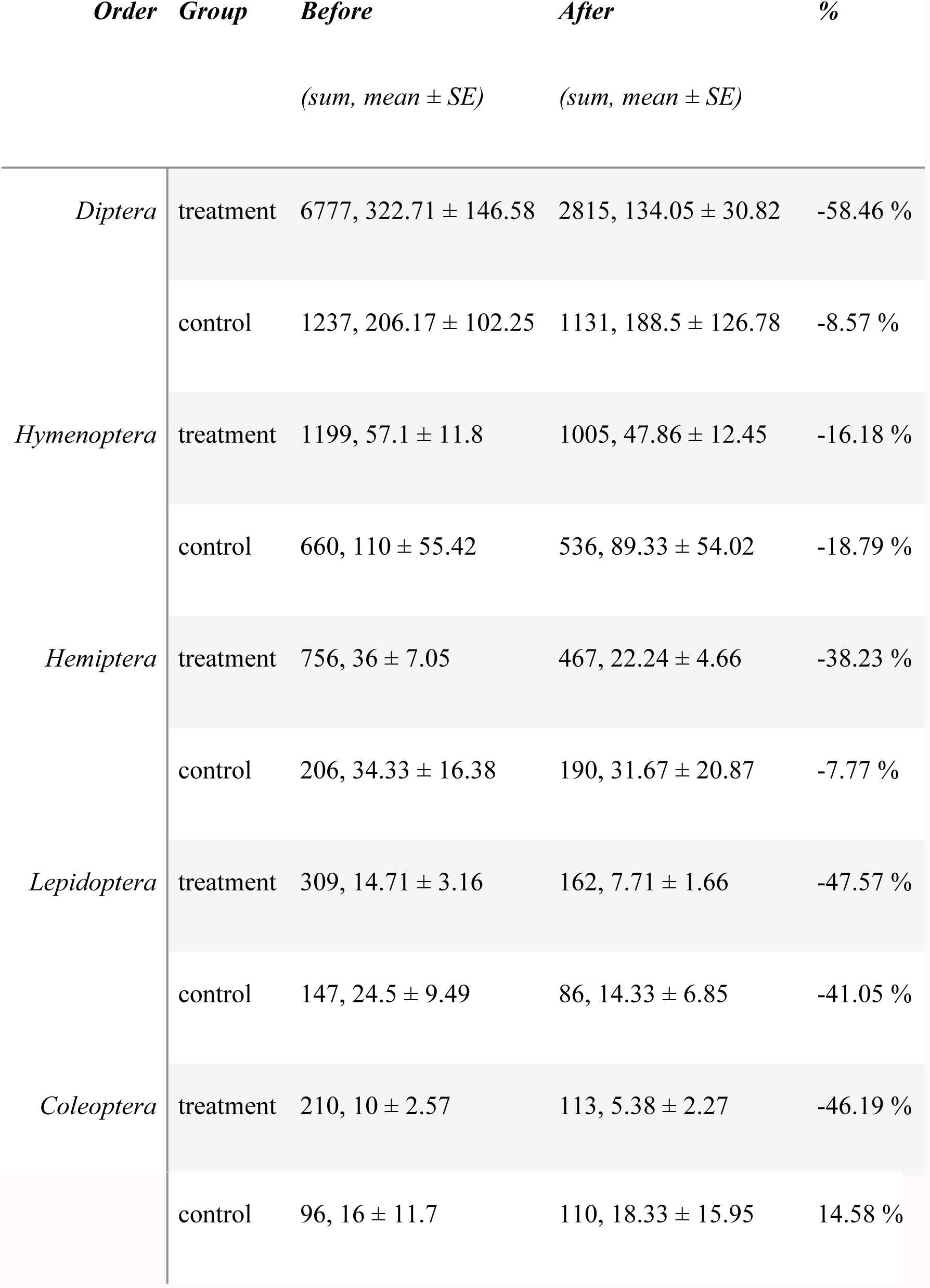
Summary statistics for the abundance (number of individuals) of different insect taxa in the malaise trap samples collected before and after insecticide spraying given separately for the treated and control sites (all sites within the same treatment group are combined).

When we treated all major pollinator taxa together (Syrphidae, Apidae and Lepidoptera), we also found that the effect of insecticide application was significant in the treatment group (group means ± SE: before treatment, 25.05 ± 4.16; after treatment, 17.38 ± 4.02; paired t-test: *t*_20_ = 3.223, *P* = 0.004, Figure 7), but not in the control group (group means ± SE: before treatment, 57.33 ± 29.14; after treatment, 46.83 ± 29.28; paired t-test: *t*_5_ = 0.805, *P* = 0.458, Figure 7).

**Figure 7.**
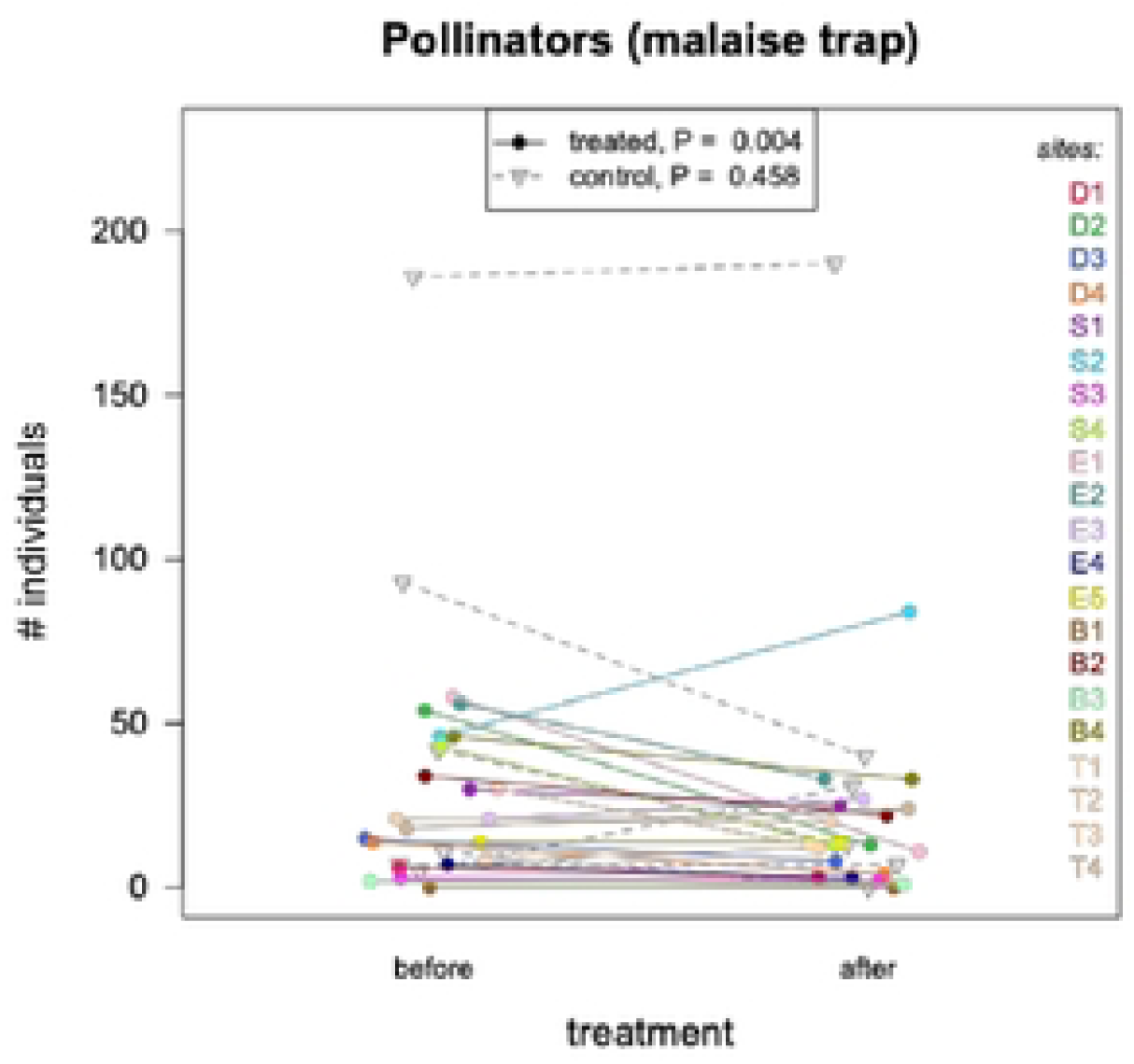
The effect of insecticide treatment on the abundance of pollinator insects detected in the malaise traps (Syrphidae, Apidae and Lepidoptera). Coloured dots are for sites where mosquito control was applied, grey triangles are for sites where such intervention was not carried out on the same date. Different colours represent different sites (with IDs shown on the right corresponding to locations shown in Figure 1), and the lines connect the detected insect abundances of the same site before and after the night of the application of insecticides. *P* values are from the corresponding paired-t test performed on the log_10_-transformed data. Points along the x-axes are jittered for better visualisation.

The main results for the non-target organisms remained unchanged in the framework based on mixed models that accounted for the hierarchical structure of data and date effects (Supplementary Material Tables S2 and S3).

## Discussion

Here, by adopting an experimental design in field conditions, we assessed how the use of ULV insecticide applications in Hungary affects the abundance of both target and non-target organisms during mosquito control. The first main result was that mosquito abundance significantly declined in the treatment group (but not in the control group) after the application of insecticides, with an approximately 45% reduction in the BG Sentinel trap catches. Second, we also showed that the effect of treatment was similar when we separated the target organisms into native and invasive pools of species, and we could uncover a significant reduction in the abundance of *Ae. albopictus* and *Ae. koreicus* following the ULV treatment. Our third key finding revealed, however, that the negative treatment effect on general mosquito abundance was not unanimously present in all sites, as in some localities the number of mosquitoes did not decrease after the day of spraying. Fourth, focusing on the non-target organisms, we also quantified how the use of ULV insecticide applications affected the community of flying insects, and demonstrated a considerable reduction of abundance of insects that are smaller than or similar in size to mosquitoes, as well as in most insect taxonomic groups, especially in pollinators.

### Effects on target organisms

Our results showed that ULV spraying produced an average reduction of more than 45% in adult mosquito abundance, which places the performance of the Hungarian ground-based control program within the middle range of reductions reported internationally (Corcos et al. 2020; Fonseca et al. 2013; Hart et al. 2024; Manica et al. 2017; Unlu et al. 2018). Previous monitoring efforts in Hungary have generally lacked rigorous experimental assessment of ULV, but available reports from other countries suggest that ULV applications often yield moderate and highly variable short-term declines in mosquito nuisance (Corcos et al. 2020; Fonseca et al. 2013; Hart et al. 2024; Manica et al. 2017; Unlu et al. 2018). Studies conducted elsewhere in Europe and North America typically document reductions of 30–80% within the first 12–24 hours following ULV adulticiding, while consistently noting that these effects are transient and strongly dependent on local environmental and operational conditions (Corcos et al. 2020; Fonseca et al. 2013), which we also discovered in the current study.

From the perspective of public health mosquito control, a reduction of less than 80% is usually considered insufficient to achieve meaningful suppression, especially in areas experiencing high nuisance levels or active arbovirus transmission (Unlu et al. 2018). However, the fact that mosquito abundance in treated sites was not statistically differentiable from the abundance of target organisms in untreated sites after spraying suggests that ULV interventions has a potential to bring population densities down to levels, which do not require insecticide treatment. For making interpretations about the efficiency of the applied mosquito control based on the content of BG Sentinel traps, one must also consider the inherent selectivity of the device (Roiz et al. 2012). These traps disproportionately attract host-seeking females of certain species while undersampling others, meaning that trap-based estimates may not fully reflect the composition and abundance of the broader wild mosquito community. Accordingly, our estimates are valid for the mosquito community represented in the trap samples but may not apply to all wild mosquito populations. In addition, we cannot formulate any conclusions regarding the lasting effect of the treatment, which is also crucial both for the perspective of public health and that of general nuisance. Our study design focused on paired short-term assessments, thus we were only able to evaluate immediate responses to the intervention, and cannot conclude whether the observed reductions persisted beyond the first sampling interval or whether rapid population rebound occurred, as documented in many other studies (Boubidi et al. 2016; Corcos et al. 2020; Fonseca et al. 2013; Manica et al. 2017; Unlu et al. 2018). Therefore, additional studies are needed that assess the changes in mosquito abundance over a broader time window and apply more diverse trapping methods to obtain more balanced samples of wild populations.

### Heterogeneity in treatment effectiveness

Despite the overall negative effect of ULV spraying on mosquito abundance, treatment outcomes varied markedly among sites, with several localities showing little or no detectable reduction following the intervention. Such heterogeneity is a common feature of ULV adulticiding and underscores the strong dependence of treatment success on local environmental and operational conditions (Boubidi et al. 2016; Corcos et al. 2020; Fonseca et al. 2013; Hart et al. 2024). Our analyses indicate that the magnitude of mosquito decline was associated with initial mosquito abundance and, independently, wind speed. Higher initial abundance likely increases the probability that host-seeking mosquitoes are active and exposed during spraying, while unfavourable wind conditions can rapidly dilute or displace insecticide droplets by its spray drift impacts (Oberhauser et al. 2009). Together, these factors can explain why identical control measures may yield pronounced effects in some locations but fail entirely in others. However, we cannot exclude the possibility that wing speed affected the efficiency of trapping – and not the efficiency of mosquito control *per se* –, as stronger winds may affect the ability of mosquitoes to fly declining their probability to enter the BG Sentinel device. Importantly, the spatial variability in the decline of mosquito abundance in the traps implies that ULV spraying cannot be assumed to provide uniform mosquito suppression across treated areas, and that suboptimal operational conditions may result in ecological exposure without achieving meaningful control of target populations. These findings highlight the need for more context-dependent planning, risk-assessment, real-time adjustment of spraying protocols, and post-treatment evaluation to ensure that ULV interventions achieve their intended outcomes.

### Effects on invasive mosquito species

When native and invasive mosquito species were analysed separately, ULV spraying resulted in a significant decline in both groups, with the proportional reduction observed for invasive mosquitoes – dominated by *Aedes albopictus* and to a lesser extent *Ae. koreicus* – being comparable to what was detected for native species. This finding is noteworthy, because invasive *Aedes* mosquitoes are often hypothesized to be poorly affected by conventional ULV adulticiding (Boubidi et al. 2016; Farajollahi et al. 2012; Fonseca et al. 2013). These expectations are grounded in the ecological and behavioural characteristics of these species, including their predominantly diurnal activity patterns, their tendency to rest in sheltered microhabitats such as dense vegetation, building structures, and private gardens, and their relatively limited dispersal distances within urban landscapes. Together, these features may reduce exposure to insecticide droplets applied during evening or night-time ground spraying operations. However, contrary to such predictions, our results demonstrate that invasive *Aedes* populations can experience short-term reductions comparable to those of native mosquitoes following ULV treatments. This suggests that a considerable proportion of the invasive mosquito populations remains active or exposed during the application period, or that insecticide droplets can penetrate vegetated or semi-sheltered urban microhabitats effectively. However, this apparent sensitivity should also be interpreted cautiously, as short-term declines do not necessarily translate into sustained population suppression, particularly for container-breeding invasive species that can rapidly replenish adult populations from untreated larval habitats (Boubidi et al. 2016; Corcos et al. 2020; Fonseca et al. 2013; Manica et al. 2017; Unlu et al. 2018). Nevertheless, our findings provide empirical field-based evidence that invasive mosquitoes in Hungary are indeed responsive to ULV spraying, and that their limited control success reported elsewhere (Faraji et al. 2016, Walker et al. 2025) may be driven more by operational and ecological constraints than by species-specific tolerance to the applied insecticides.

### Effects on non-target organisms

Beyond its effects on mosquitoes, ULV spraying had a pronounced impact on non-target flying insects, as revealed by the malaise trap samples. At treatment sites, the abundance of non-target insects declined by more than 40% on average, a reduction comparable in magnitude to that observed for mosquitoes. This effect was consistently detected across most sites and was evident in multiple major insect orders, including Diptera (excluding mosquitoes), Hemiptera, Lepidoptera, and Coleoptera. Importantly, pollinators depicted a considerable decline following the treatment. When insects were grouped by body size, the strongest negative effects were observed among small- and medium-sized taxa – those most similar to mosquitoes in size – whereas large-bodied insects showed no statistically detectable decline. This size-dependent pattern is consistent with the physical properties of ULV droplets and with previous studies demonstrating that larger flying insects are less likely to intercept insecticide aerosols during spraying events or they are less likely to die from the same doses of ingredients due to their larger body size (Boyce et al. 2007; Corcos et al. 2020; Kwan et al. 2009). The widespread reduction across taxonomic groups indicates that ULV applications are not selective for mosquitoes, but affect a broad spectrum of aerially active insects, many of which contribute to essential ecosystem functions such as pollination (Syrphidae, Apidae and Lepidoptera). Although the magnitude of decline varied among taxa, the overall pattern provides strong field-based evidence that ground-based ULV spraying imposes substantial short-term ecological costs on non-target insect communities, reinforcing concerns raised by earlier experimental and observational studies from other regions (Boyce et al. 2007, Davis & Peterson 2008, Oberhauser et al. 2009, Bonds 2012, Abeyasuriya et al. 2017, Bargar & Jiang 2024). However, the shortcoming that arises from the short temporal window of the study design also applies to the sampling of the non-target insect communities, thus we cannot make any conclusion about the long-lasting ecological side-effects of ULV treatment.

### Study limitations

Several limitations of this study should be acknowledged when interpreting the findings. First, although the paired before–after control–treatment design strengthens causal inference, the overall sample size was necessarily limited by the availability of suitable sites and coordinated spraying events, which constrained the complexity of statistical models and the number of covariates that could be evaluated simultaneously. Second, our assessment relied on trap-based sampling, which inherently reflects only a subset of the mosquito and insect communities present. In particular, BG Sentinel traps selectively attract host-seeking females of certain mosquito species and may underrepresent taxa with different activity patterns or sensory preferences, while malaise traps also have their specific sensitivity to specific insect groups and behaviours. Third, the temporal scope of the study was restricted to short-term responses immediately following ULV application, preventing assessment of population rebound, delayed mortality, or longer-term ecological recovery of non-target insect communities. Fourth, although we investigated several environmental variables associated with treatment effectiveness, these factors were not experimentally manipulated and may covary in complex ways that cannot be fully disentangled in an observational field study. Finally, our analyses focused on changes in abundance rather than on demographic or fitness-related parameters, such as survival, reproduction, or community composition over time. Future research should therefore incorporate longer monitoring periods, repeated treatment events, and a broader range of ecological indicators, ideally with comparison with alternative vector management approaches, to more fully evaluate the efficacy, sustainability, and ecological consequences of mosquito control practices.

## Conclusion

Taken together, our results demonstrate that ground-based ULV insecticide applications in Hungary provide considerable short-term reductions in mosquito abundance with both native and invasive mosquito species depicting measurable declines following treatment. However, our analysis yielded inconsistencies in effectiveness among sites pointing to the sensitivity of ULV treatment to logistical and environmental conditions. Furthermore, we also demonstrated that the reduction of the abundance of mosquitoes was accompanied by the substantial negative effects on non-target flying insect communities involving multiple taxonomic groups, highlighting a poor selectivity of the applied control method. This imbalance between the transient benefits and the ecological costs raises concerns about the sustainability of ULV spraying as a primary mosquito control strategy. Our findings underscore the need to re-evaluate current mosquito management practices in Hungary and similar regions, and to shift toward more ecologically informed approaches that prioritize larval source reduction, biological control, and targeted interventions within an integrated vector management framework. Incorporating systematic post-treatment evaluation and ecological impact assessments into control programs along the design we followed in this study would be essential to ensure that public health objectives are met without imposing disproportionate harm on insect biodiversity and ecosystem functioning.

## Acknowledgement

We thank Gábor Fekete (Corax-Bioner Co.) and Zoltán Kenyeres (research group of Pannónia Központ Ltd) for providing detailed information about the mosquito control program and for their cooperation throughout this study. We are also grateful to all volunteers who participated in the sample collection. The study was supported by funds from Hungary’s National Research, Development and Innovation Office (K135841, RRF-2.3.1-21-2022-00006, ADVANCED 152427) and by the AXA Research Fund. G.M. was supported by the Research Excellence Programme of the Hungarian University of Agriculture and Life Sciences (KKP2024-MG, KKP2026-MG). Z.S was supported by the National Research, Development and Innovation Office (FK-147466). The acknowledged parties had no role in study design, data collection and analysis, decision to publish, or preparation of the manuscript.

## Supplementary Material

**Table S1.**
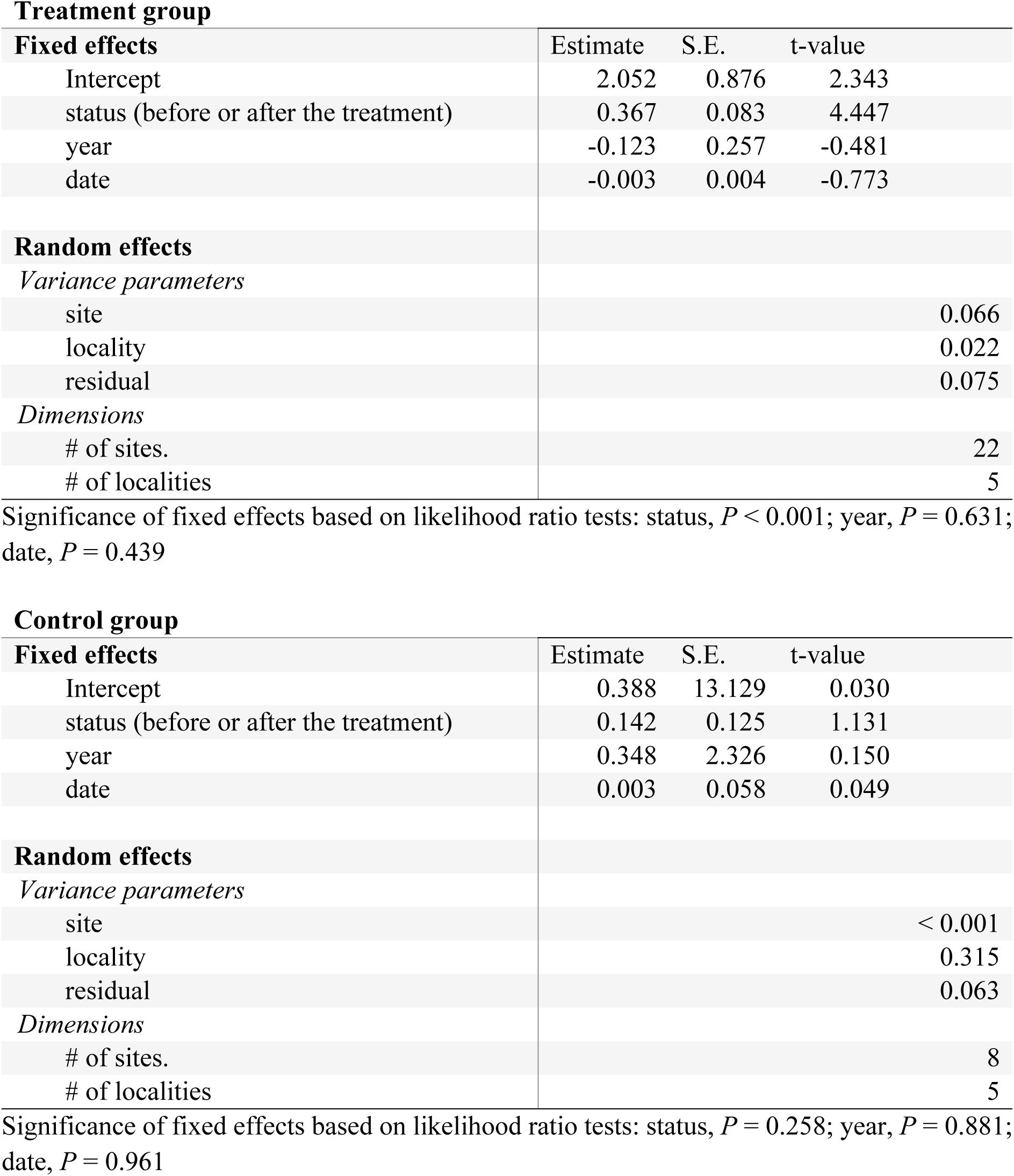
Results of the mixed models testing for the effect of treatment on mosquito abundance (number of individuals) when accounting for the hierarchical structure of data and year and date effects.

**Table S2.**
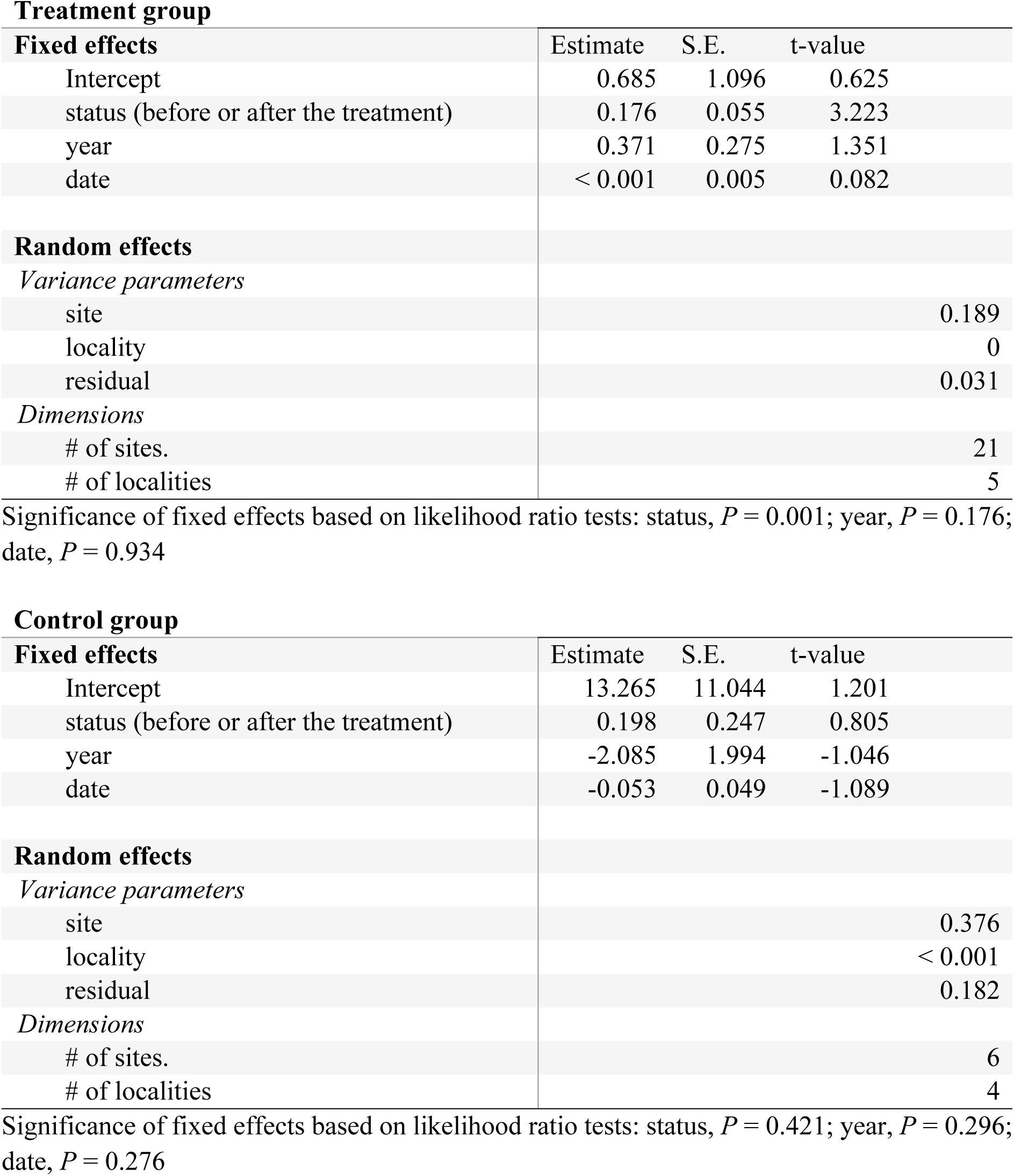
Results of the mixed models testing for the effect of treatment on the abundance of pollinators when accounting for the hierarchical structure of data and year and date effects.

**Table S3.** Results of the mixed models testing for the effect of treatment on the abundance of all insects that were captured in the malaise traps when accounting for the hierarchical structure of data and year and date effects.

**Treatment group**
| **Fixed effects** | Estimate | S.E. | t-value |
| --- | --- | --- | --- |
| Intercept | 3.383 | 1.313 | 2.576 |
| status (before or after the treatment) | 0.190 | 0.060 | 3.158 |
| year | 0.396 | 0.332 | 1.193 |
| date | -0.007 | 0.006 | -1.350 |
| **Random effects** |  |  |  |
| *Variance parameters* |  |  |  |
| site |  |  | 0.252 |
| locality |  |  | 0.007 |
| residual |  |  | 0.038 |
| *Dimensions* |  |  |  |
| # of sites. |  |  | 21 |
| # of localities |  |  | 5 |

**Control group**
| **Fixed effects** | Estimate | S.E. | t-value |
| --- | --- | --- | --- |
| Intercept | 12.516 | 9.999 | 1.252 |
| status (before or after the treatment) | 0.259 | 0.132 | 1.960 |
| year | -1.754 | 1.805 | -0.972 |
| date | -0.046 | 0.044 | -1.044 |
| **Random effects** |  |  |  |
| *Variance parameters* |  |  |  |
| site |  |  | 0.357 |
| locality |  |  | < 0.001 |
| residual |  |  | 0.052 |
| *Dimensions* |  |  |  |
| # of sites. |  |  | 6 |
| # of localities |  |  | 4 |

Significance of fixed effects based on likelihood ratio tests: status, $P = 0.050$ ; year, $P = 0.331$ ; date, $P = 0.297$

**Table S4.**
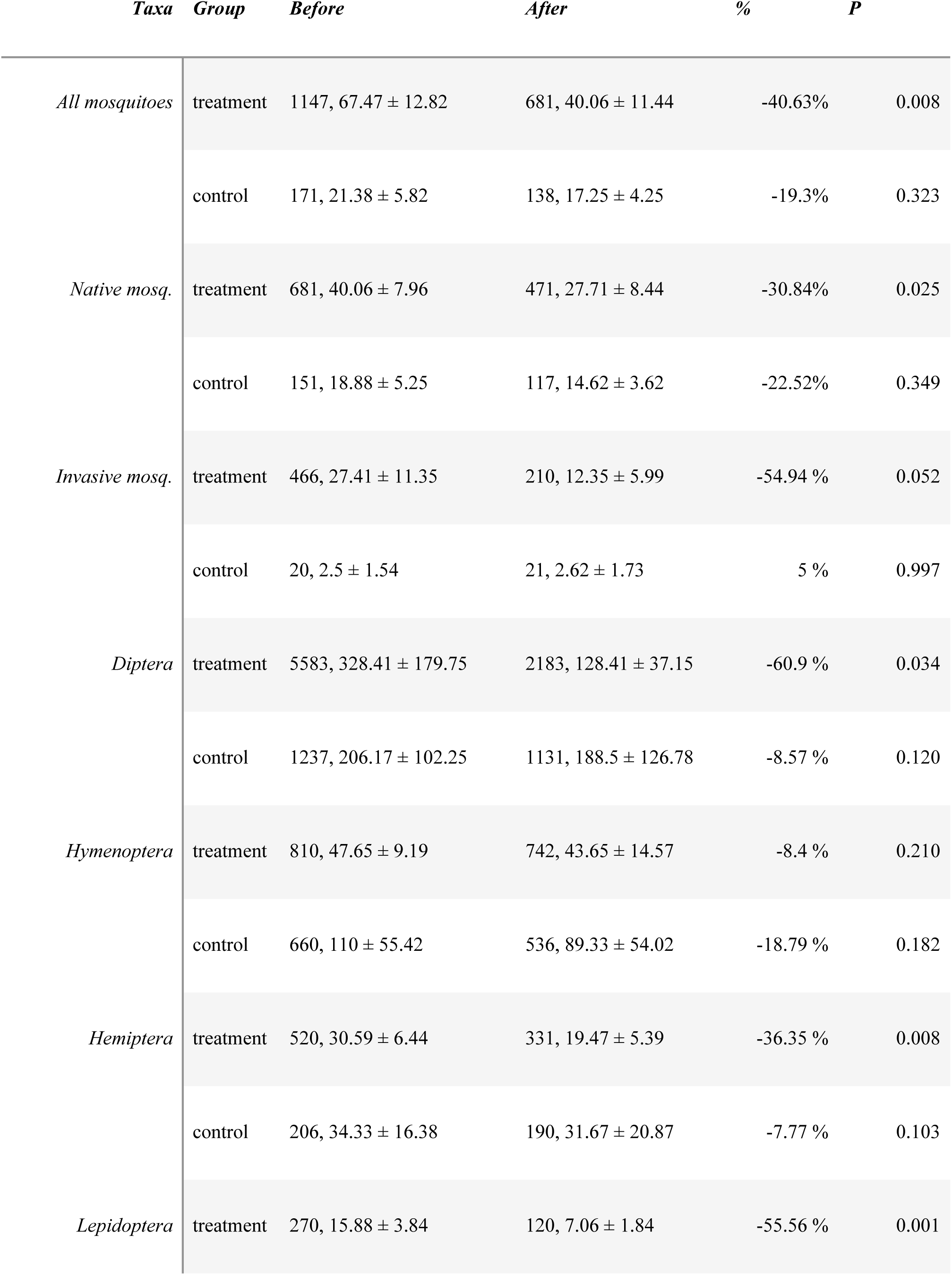
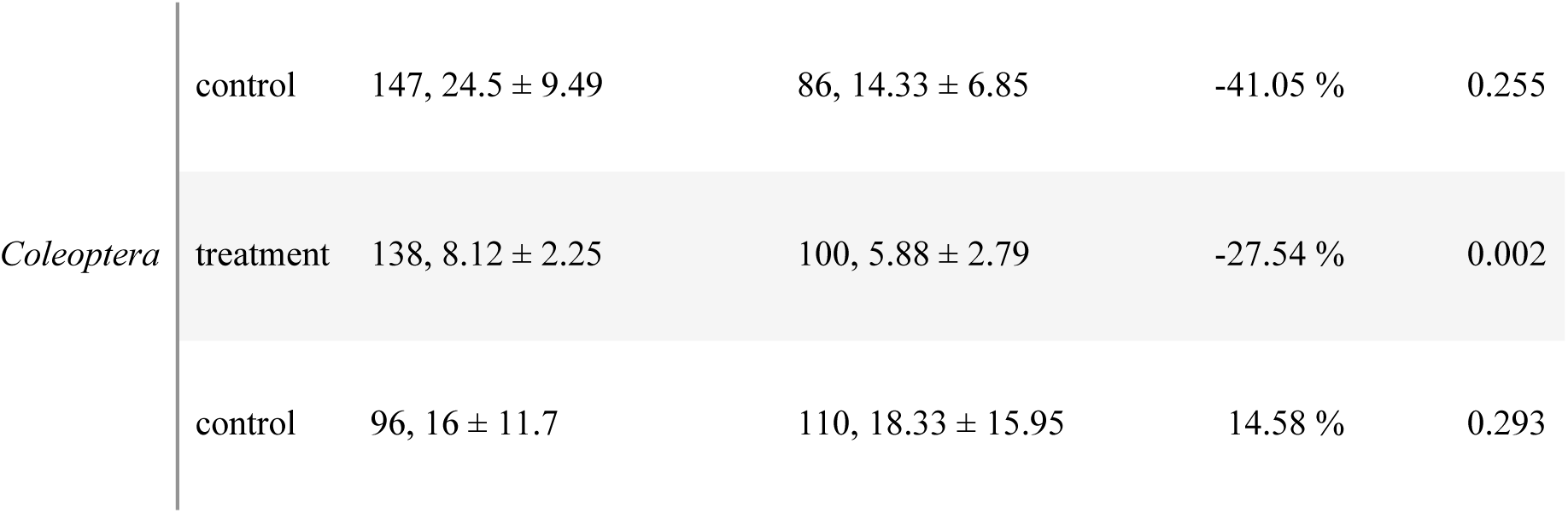
Summary statistics for the abundance (number of individuals) of different insect taxa in the malaise trap samples collected before and after insecticide spraying given separately for the treated and control sites (all sites within the same treatment group are combined) after excluding catches from Tiszafüred. *P* values are for the corresponding paired t-tests

